# A deep learning classification task for brain navigation during functional ultrasound imaging

**DOI:** 10.1101/2022.03.18.484747

**Authors:** Théo Lambert, Clément Brunner, Dries Kil, Roel Wuyts, Ellie D’Hondt, Gabriel Montaldo, Alan Urban

## Abstract

Positioning and navigation are essential components of neuroimaging as they improve the quality and reliability of data acquisition, leading to advances in diagnosis, treatment outcomes, and fundamental understanding of the brain. Functional ultrasound (fUS) imaging is an emerging technology providing high-resolution images of the brain vasculature, allowing for the monitoring of brain activity. However, as the technology is relatively new, there is no standardized tool for inferring the position in the brain from the vascular images. This study presents a deep learning-based framework designed to address this challenge. Our approach uses an image classification task coupled with a regression on the resulting probabilities to determine the position of a single image. We conducted experiments using a dataset of 51 rat brain scans to evaluate its performance. The training positions were extracted at intervals of 375 µm, resulting in a positioning error of 176 µm. Further GradCAM analysis revealed that the predictions were primarily driven by subcortical vascular structures. Finally, we assessed the robustness of our method in a cortical stroke where the brain vasculature is severely impaired. Remarkably, no specific increase in the number of misclassifications was observed, confirming the method’s reliability in challenging conditions. Overall, our framework provides accurate and flexible positioning, not relying on a pre-registered reference but on conserved vascular patterns.

## Introduction

Matching brain images with a reference, a process called registration is essential for both medical and research purposes. In clinical applications, it is used to facilitate diagnosis and real-time navigation during neurosurgery^1–4^. In preclinical research, brain images are often registered to an atlas (e.g., the Allen^5^ or Paxinos^6^ atlas for rodents) to identify anatomical regions and interpret the data. In this context, each imaging modality requires specific registration tools that must be co-developed with the imaging modality^7–12^.

Applied in both preclinical (e.g., rodents^13–16^ and non-human primates^17,18^) and clinical contexts (e.g., neonates^19,20^ and adults^21–23^), functional ultrasound (fUS) imaging is a recently established technology that combines large depth-of-field and high spatiotemporal resolution^24–26^. By repeating the acquisition of micro-Doppler images over time, fUS imaging tracks the hemodynamics of small vessels^13,24,27^, reflecting neuronal activity^15,28–31^.

Manual registration of micro-Doppler images is typically performed by matching landmarks (e.g., cortical surface, major brain regions, and vessels) with a reference atlas^32^. Performing the precise localization of single images with this process is challenging as it does not fully account for morphological variability across individuals or models^33,34^. Furthermore, this approach is time-consuming and subject to inter-operator variability. Therefore, there is a strong need for a positioning system that accurately infers anatomical positions from the conserved brain vasculature^35–37^.

An automated methodology has recently been proposed for ultrasound-based neuro-navigation on mice^38^. It relies on the online registration of a micro-Doppler scan, acquired at the beginning of every experiment, to a reference micro-Doppler volume pre-registered to an atlas. Yet, this initial brain scan makes the approach incompatible with all experimental contexts, such as small imaging windows and larger brains where only a limited portion is imaged. Moreover, relying on an atlas for positioning does not, in essence, fully account for morphological variability across individuals or models^33,34^.

We designed a deep learning-based classification framework leveraging convolutional neural networks (CNN) to circumvent these limitations. Solely based on the brain vasculature, our approach allows the localization of single micro-Doppler images with 176µm precision for both positioning and navigation applications. This strategy provides an accurate, reference-free, and user-independence that can be extended to various experimental designs and models, from larger animals up to human applications.

## Results

### Operating principle

Our method uses a neural network trained on a classification task to find the position of a given image. Three main elements are required for the implementation: 1) an input dataset of images aligned to a predefined reference, 2) a neural network trained to classify images depending on their position, and 3) a probability-based regression that infers the position of a new input image. These parts are detailed below in the context of rat brains and with a DenseNet121-CNN. Nevertheless, it is essential to note that this operating principle is not species-nor neural network-specific.

#### 1) Input dataset

In this study, we used a set of n=51 rat brains. Each brain was scanned with 77 micro-Doppler images extending from Bregma +3.0 to -6.5 mm with an in-between image spacing of 125 µm (**Fig. 1-a**). A micro-Doppler image has an in-plane resolution of 100×110 µm and 300-µm slice thickness in this setting^13,24,32^. Example micro-Doppler images on which major anatomical structures and vessels were annotated are shown in **Fig. 1-b**, and **Supplementary Fig. S1**. Scans were aligned on reference positions by expert consensus (**Materials and Methods – Registration of micro-Doppler images**).

**Fig. 1:**
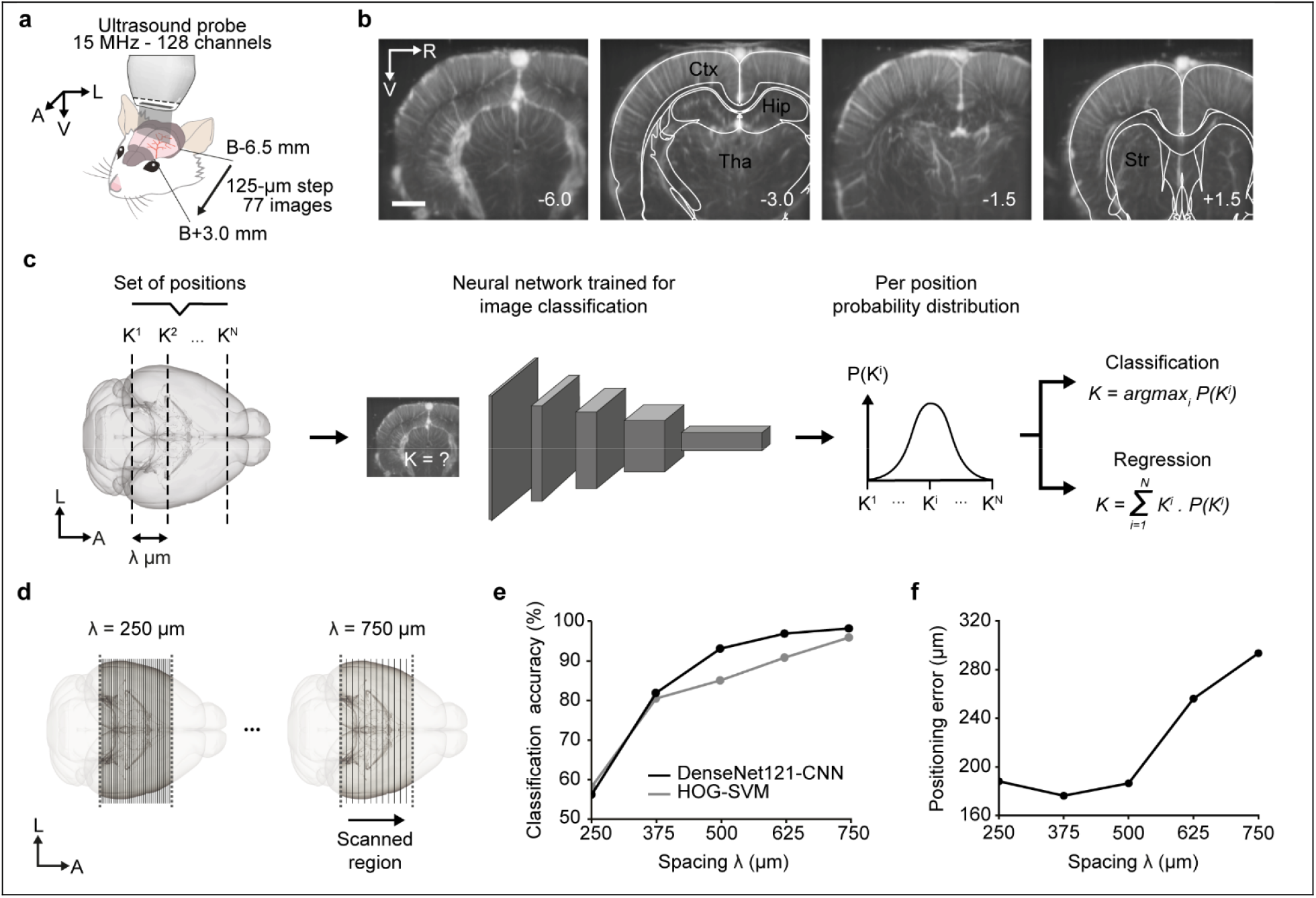
Methodological approach and spacing selection. **a-**Acquisition setup for large-scale micro-Doppler imaging of rat brains. The ultrasonic probe is moved along the postero-anterior axis using a motorized linear stage. The imaging was performed from Bregma -6.5 to +3.0 mm with a 125 µm spacing for a total number of 89 images. **b-**Set of micro-Doppler images extracted from a single scan overlaid with a simplified version of the Paxinos brain atlas^6^ in white. Main anatomical structures are identified in black: Cortex (Ctx), Hippocampus (Hip), Thalamus (Tha), and Striatum (Str). The Bregma position (in mm) of the micro-Doppler images is shown in the lower right corner. Scale bar: 2 mm. **c-**Schematic representation of the position inference procedure. Left: A set of positions (K^1^, …, K^N^) with spacing λ µm is defined over the brain. Center: An image with unknown position K is fed to a neural network trained to classify input microDoppler images depending on their position. Right: the network outputs a probability distribution over the positions (K^1^, …, K^N^). It can be used either to determine K as being the most likely position within (K^1^, …, K^N^) (classification) or to estimate K as a weighted sum of all positions using their respective probabilities (regression). **d-**Down-sampling procedure used to determine an optimal set of positions. Each scan is down-sampled using 5 factors, corresponding to an increase in the spacing λ between two consecutive images: 250, 375, 500, 625, and 750 µm. **e-**Classification accuracy (%) of DenseNet121-CNN (black) and HOG-SVM (grey) models for each spacing (testing, n=13 rats). **f-**Regression error (µm) obtained from the DenseNet121-CNN model for each spacing (testing, n=13 rats). A: anterior, L: left, R: right, V: ventral.

#### 2) Neural network-based image classification

For the needs of this study, we constrained the selection to classical models. After a preliminary performance evaluation (**Supplementary Table S1, Materials and Methods – Model selection**), a DenseNet121-CNN and HOG-SVM with additive χ^2^-kernel were selected as main and baseline models, respectively^39,40^. Both were trained to classify input micro-Doppler images to their position on a subset of 25 rats for a given set of positions (**Fig. 1-c)**. The hyperparameters were tuned on a validation set of 13 rats, and their performance was assessed on a testing set of 13 rats.

#### 3) Navigation stage

Given a new image with an unknown position, the CNN outputs a probability for each position included in the training dataset. The training positions themselves are used as regressors, and their probabilities as corresponding coefficients in a regression model, providing the actual position estimate (**Fig. 1-c**).

To assess the performance of our approach, we evaluated the accuracy of the classifier and the associated positioning error, analyzed their spatial dependence, and extracted the anatomical regions supporting the inference.

### Effect of the scanning spacing in the input dataset

To determine a suitable spacing to scan the brains in the input dataset, we created five datasets by down-sampling the original brain scans at 250 µm (39 positions), 375 µm (26 positions), 500 µm (20 positions), 625 µm (15 positions) and 750 µm (13 positions) (**Fig. 1-d**).

Both DenseNet121-CNN and HOG-SVM show an increase in the classification accuracy along with the spacing (250 to 750 µm), from 56.2% to 98.2% and from 58% to 95.9%, respectively (**Fig. 1-e** and **Supplementary Table S2**). The training and validation accuracies follow similar trends, indicating that the models did not overfit training data in a problematic way (**Supplementary Fig. S2**). As an additional control, we performed a 5-fold cross-validation on the 375 µm dataset, which resulted in comparable testing accuracy (80.9 ± 2.3%). It should be noted that the CNN did not converge with 125-µm spacing.

The testing accuracy is lower for the HOG-SVM than the DenseNet121-CNN, irrespective of the spacing. However, the performance comparison using the McNemar^41^statistical test exhibited no statistically significant differences from spacings 500 and 625 µm (^**^p=0.0012 and ^*^p=0.012, respectively; **Supplementary Table S2**). Regarding the extrema, both models achieve similar maximum class accuracy, while the DenseNet121-CNN provides higher and lower minimum class accuracy than the HOG-SVM for spacings 625 / 500 µm and 375 / 250 µm *respectively*.

Although better for the classifier accuracy, increasing the spacing decreases the resolution of the positioning. To determine the value that gives the most accurate results, we estimated the positioning error. It is defined as the standard deviation of the differences between the predicted positions and the target positions in the animals of the validation set (**Fig. 1-c, Materials and Methods – Probability-based regression**). The 375 µm spacing dataset gave the best result with a regression error of 176 µm and was therefore selected for further investigation. For comparison purposes, we also computed the positioning error with the other CNN models (i.e., ResNet50, DenseNet121, VGG11, ViT-Base-16, and EfficientNetV2-S; see **Supplementary Table S1**).

### Spatial dependency of the classification

To assess the reliability of the inference in different parts of the brain, we examined the classification accuracies by position on the dataset with 375µm spacing. The values are non-uniformly distributed and range from 30.8 to 100% (**Fig. 2-a**). The DenseNet121-CNN and HOG-SVM displayed similar results overall, and the anterior part of the brain generally elicited lower accuracies for both models (5 of the 8 positions with accuracy below the mean are between Bregma 0.0 to +3.0 mm). The DenseNet121-CNN-associated confusion matrix reveals that misclassified images were mapped to neighboring positions (**Fig. 2-b**) and were not concentrated in a subset of rats.

**Fig. 2:**
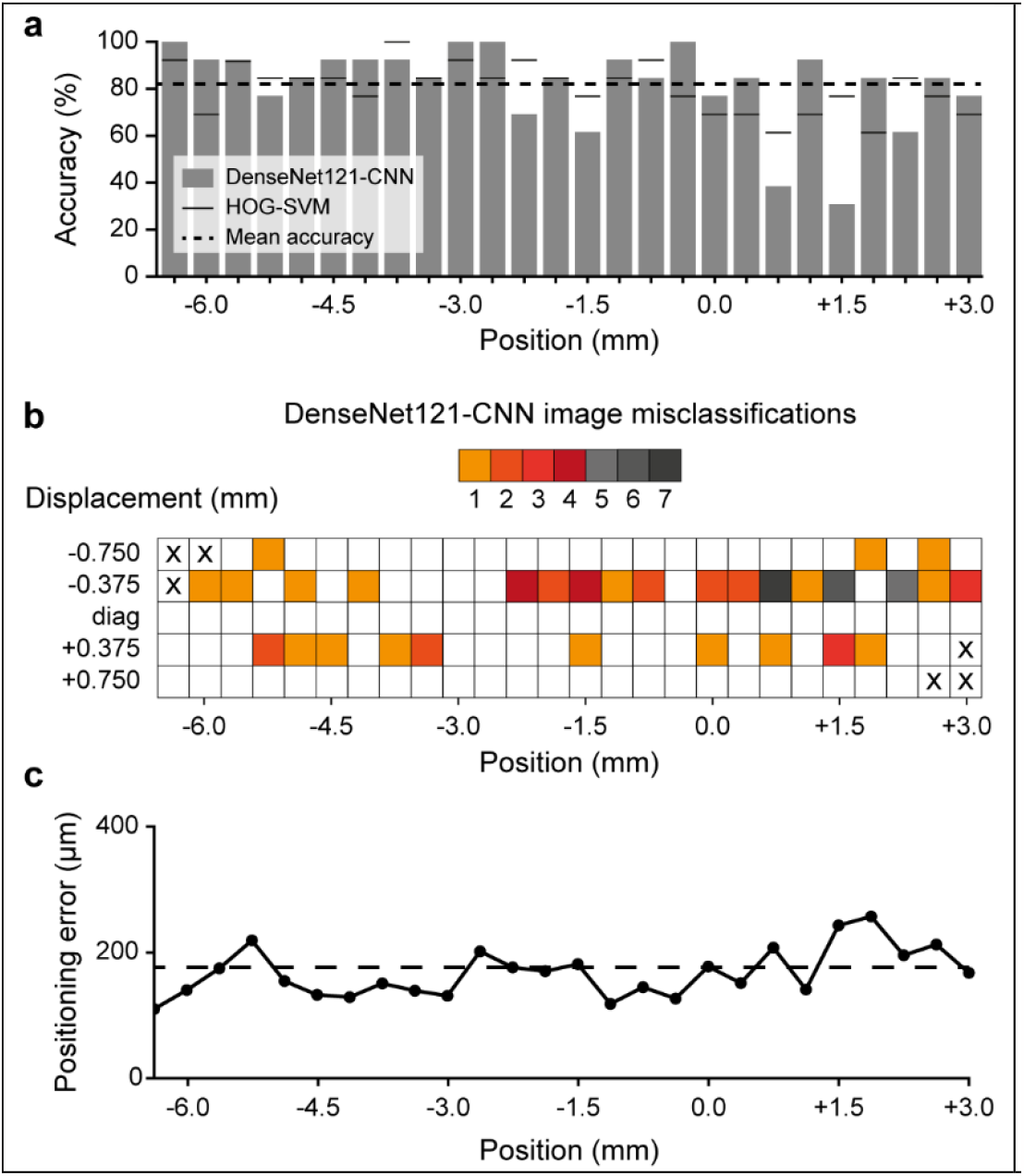
Per position analysis of the DenseNet121-CNN predictions. **a-**Per position display of the DenseNet121-CNN accuracies for the dataset with 375-µm spacing (testing, n=13 rats). The horizontal dashed line represents the mean classification accuracy. For each position, the black dash represents the accuracy of the HOG-SVM model. **b-**Representation of the extended diagonal of the DenseNet121-CNN confusion matrix. Each arrow goes from the correct position towards the predicted position. The different colors represent the number of misclassified micro-Doppler images (375 µm spacing, testing, n=13 rats). **c-**Distribution of positioning error to the Bregma position (DenseNet121-CNN trained on 375 µm spacing dataset, testing, n=13 rats).

We then computed the positioning error per position for the DenseNet121-CNN model (**Fig 2.c, Materials and Methods – Probability-based regression**). Although not fully proportional, the distribution of positioning errors closely matches the classification accuracies, especially with higher errors in the anterior part of the brain.

### Spatial localization of the discriminative patterns

We computed the gradient-weighted class activation maps (GradCAM)^42^ to visualize the discriminative features underlying the accurate classification. This visualization technique highlights the area of an image contributing the most to the network’s inference of a given prediction. We registered the 2D scans with a digital version of the rat Paxinos atlas^6^ (as previously done by Brunner *et al*.^43^) to i) adjust for potential differences in probe positioning and ii) allow for inter-animal comparison. For each position, the GradCAM results were averaged across animals, and a threshold was applied to mitigate the effect of interpolating such low-resolution heatmaps (**Fig. 3-a**).

**Fig. 3:**
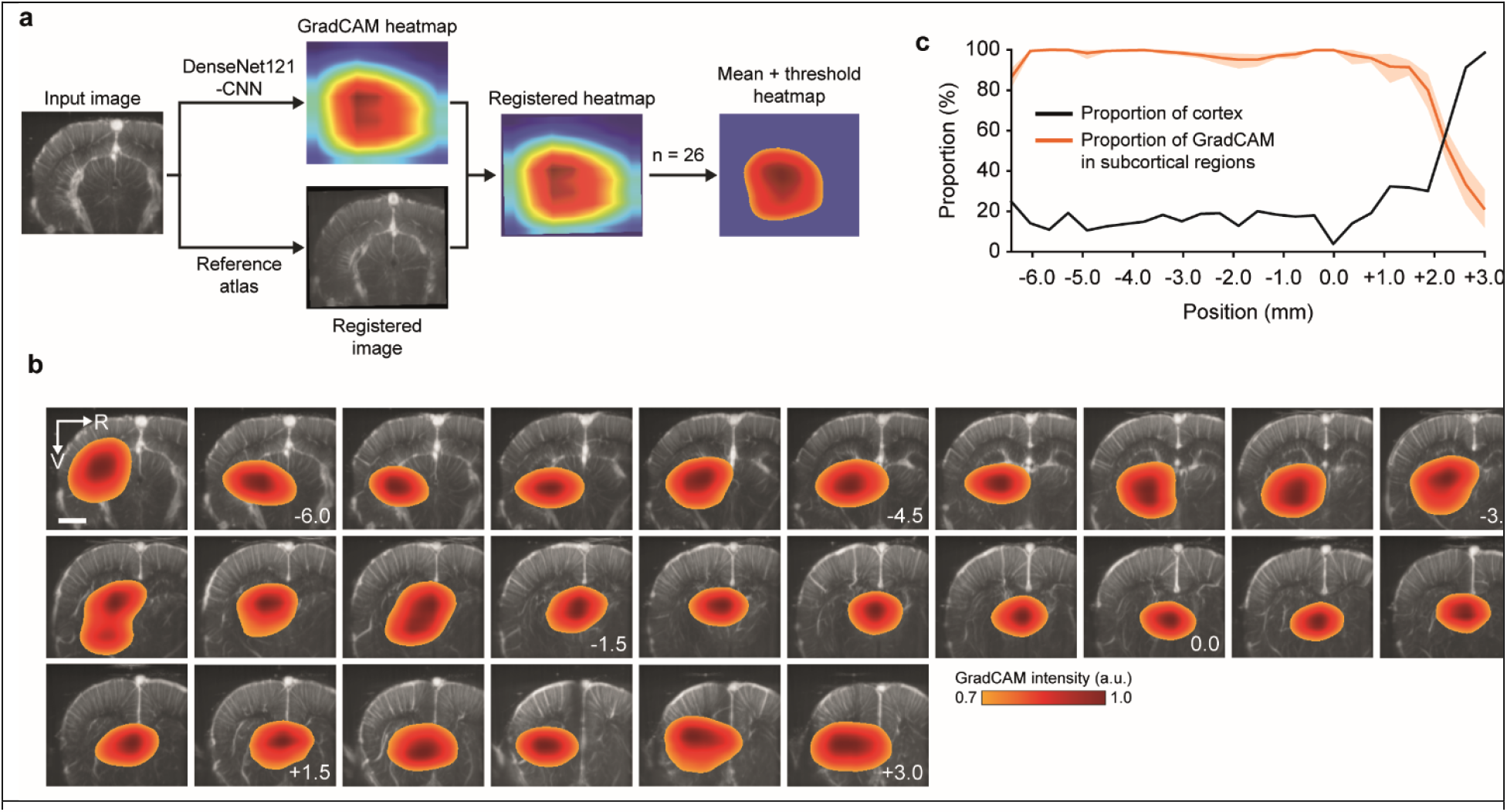
Visualization of the predictions using Gradient-weighted Class Activation Map (GradCAM) **a-**Workflow for generating the average GradCAM heatmap. Each image is processed through the DenseNet121-CNN model to collect the GradCAM heatmaps and is registered in parallel for in-plane alignment. The 26 heatmaps, including validation and testing sets, are averaged, and a threshold of 0.7 is applied. **b-**Averaged GradCAM heatmaps overlaid on corresponding (registered) micro-Doppler images. The color scale indicates the GradCAM intensity (a.u., arbitrary unit). The Bregma position (in mm) of the micro-Doppler images is shown in the lower right corner. Scale bar: 2 mm. **c-**Proportion of the GradCAM heatmap in subcortical regions (orange). The error band corresponds to the 95% confidence interval. The black curve displays the reference proportion of cortex for each position.

The average maps were overlaid per position on the corresponding registered micro-Doppler images (**Fig. 3-b**). Interestingly, a single part of the micro-Doppler image is driving the classification regardless of the location in the brain, except for Bregma -2.625 and -1.875 mm exhibiting two small, connected areas. Furthermore, the GradCAM heatmaps are almost entirely located in the subcortex up to Bregma, where the proportion starts to decrease in favor of the cortex (**Fig. 3-c**). It corresponds to the general increase in the proportion of cortex in the image.

As the GradCAM maps appeared to be conserved across rats, we identified the associated local brain vasculature. It revealed that several branches of large vessels play a significant role in the classification process. Most of these vessels supply brain regions located in subcortical regions, such as the thalamus, the hippocampus, and the striatum^35,36^, as shown in **Fig. 1-b, Fig. 3-c**, and **Supplementary Fig. S1**.

Specifically, the most important vessels in the classification of the posterior part (Bregma -6.375 to 0.0 mm) include the thalamo-perforating arteries branching from the posterior cerebral artery (PCA), the thalamostriate veins and branches (tlv), and the patterns created by adjacent vessels such as the great cerebral vein of Galen (GcvG) and the longitudinal hippocampal veins (lhv). The classification of the anterior part (Bregma 0.0 to +3.0 mm) mostly relies on the anterior cerebral artery (ACA), the azygos pericallosal arteries (APCA), and the thalamostriate veins/arteries.

### Model robustness evaluation on a cortical stroke model

To further validate the reliability of the highlighted subcortical vascular patterns and the robustness to pathological conditions, we assessed the performance of the DenseNet121-CNN on a cortical stroke model. From the 51 rats of the input dataset, a subset of 26 rats were subjected to cortical stroke by means of the permanent occlusion of the left middle cerebral artery (MCA) distal branch, provoking a significant decrease of signal (-60%) in the cortex of the left hemisphere. Brain-wide micro-Doppler scans were acquired after stroke onset (top and bottom row, respectively; **Fig. 4-a**). More details about the experimental procedure and quantification are available in ref.^43^. It should be noted that the anterior positions are minimally affected by the signal loss because those territories are poorly supplied by the MCA^36^.

**Fig. 4:**
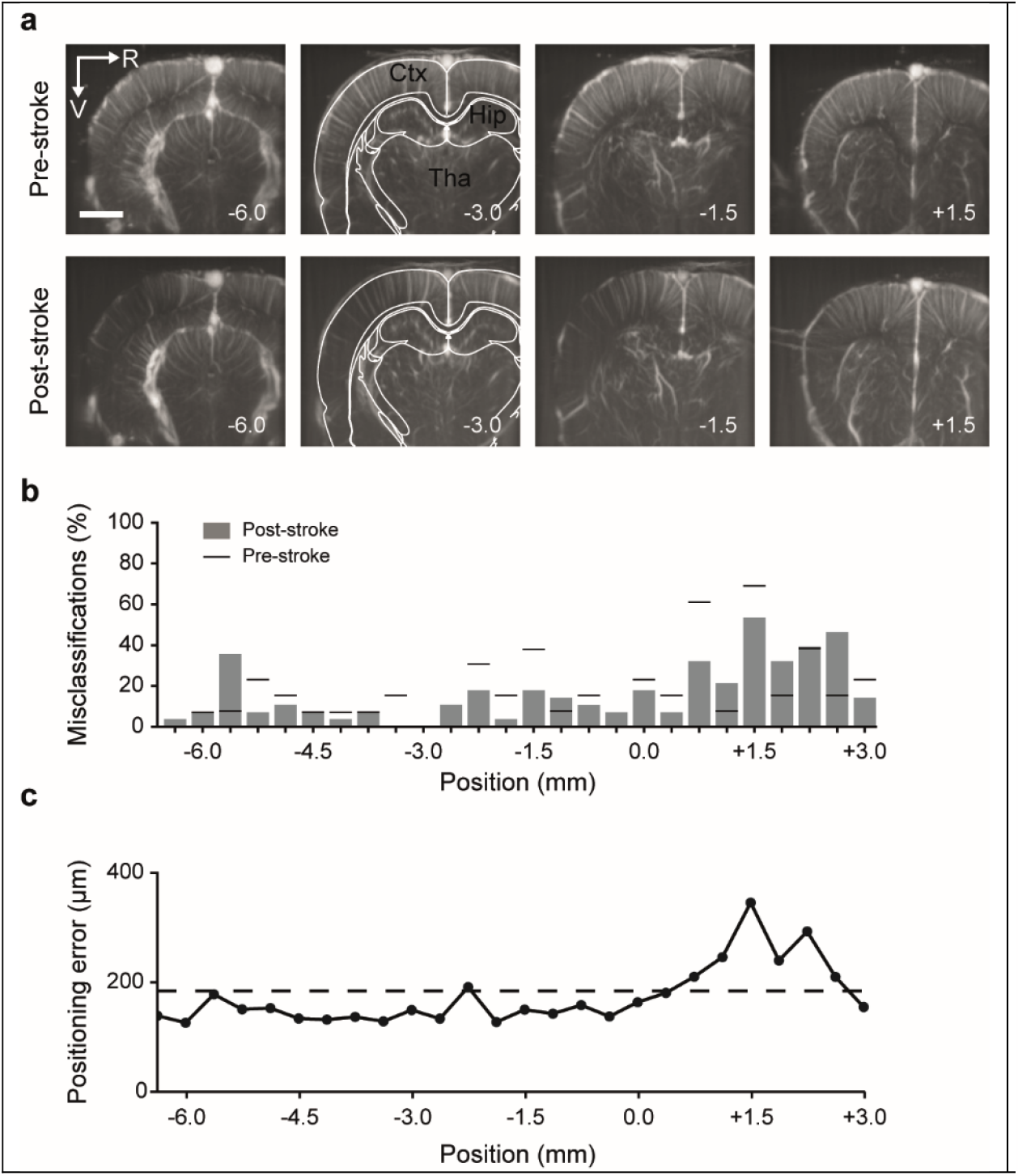
Effect of a cortical stroke on the classification accuracy. **a-**Set of micro-Doppler images before (Pre-stroke -top row) and after stroke induction (Post-stroke -bottom row). A simplified version of the Paxinos atlas^6^ is overlaid in white. Main anatomical structures are identified in black: Cortex (Ctx), Hippocampus (Hip), and Thalamus (Tha). The position from Bregma (in mm) of the micro-Doppler images is shown in the lower right corner. Scale bar: 2 mm. **b-**Proportion of micro-Doppler images misclassified before (Pre-stroke, black dash) and after (Post-stroke, grey bar) the stroke induction at each position for DenseNet121-CNN (n=26 rats). **c-**Distribution of positioning error to the Bregma position after stroke induction (DenseNet121-CNN trained on 375 µm spacing dataset, n=26 rats).

The positions of the post-stroke images were predicted without prior retraining, and the proportion of post-stroke misclassifications was computed. Images of animals originally present in the training, validation, and testing sets were pooled for this experiment. Overall, 16.5% of the images were misclassified compared to the 18.1% on the pre-stroke test set. Looking at the distribution of misclassifications per position, pre- and post-stroke exhibit similar results with only local variations (**Fig. 4-b**). Likewise, the regression error did not display significant post-stroke change in average error (176 vs. 185 µm, respectively pre/post), with consistently higher positioning error from Bregma (**Fig. 4-c**).

## Discussion

In this study, we proposed a classification framework suited for accurate and robust brain positioning and neuro-navigation during fUS imaging. Our approach relies on a neural network-based image classification task to identify a set of training positions. It serves as a reference frame from which the location of a micro-Doppler image can be inferred with precision through a regression. We have selected a DenseNet121-CNN as the main model and a HOG-SVM as the baseline model; nevertheless, the methodology is not neural network-specific and can be applied to other architectures.

First, we have defined a set of anatomical reference positions that act as classes in our classification task. By analyzing the effect of differently spaced positions, we concluded that 375 µm provides an optimal positioning error of 176 µm. We computed the positioning errors with the other models included in the model selection process. ResNet50 and VGG11-CNN yielded worse results (∼300µm positioning error), while ViT-Base-16 and EfficientNetV2-S-CNN gave errors comparable to DenseNet121, thus demonstrating the framework’s versatility.

As a result, the position of an unknown image can be determined with an error of 352 µm with 95% confidence. It is consistent with the size of the micro-Doppler image thickness (∼300 µm). Such 375-µm spacing is a trade-off between the resolution and the classification confidence: indeed, using a larger spacing allows more accurate classification but at the cost of less precision in inferring the final position. Conversely, a small spacing that is too small has a detrimental effect on the overall positioning error, which can be attributed to the vascular similarity in adjacent planes when exceeding the technology resolution.

Further analysis using the GradCAM visualization technique revealed that the classification was mainly driven by highly consistent vascular structures in subcortical regions. Since ImageNet pretraining is known to introduce a bias towards texture differences^44^, the richer vascular texture of the subcortex may account for its prevalence compared to the cortex. It can also explain why the cortical curvature and thickness variation across anatomical locations are not essential in our approach.

Finally, we validated the CNN’s predictions in rats subjected to cortical stroke, which can be considered a challenging re-test experiment. The analysis revealed no significant differences in the number of misclassifications, except for local variations, thus confirming the robustness and reproducibility of the inference in challenging contexts.

Nevertheless, it should be noted that anterior positions in our datasets exhibited higher positioning errors following the stroke induction while not being the most affected areas^43^. This observation further highlights the difficulty of performing accurate positioning anterior to Bregma +1.0 mm. It can be partially attributed to the similarity of the vascular patterns across positions in this area (cortical vessels, anterior cerebral artery, and thalamo-perforating vessels). To circumvent this issue, a first strategy could be to perform the positioning using a reference located farther away (e.g., before Bregma +1.0 or +3.0 mm) at the cost of a potentially wider cranial window. Another way could be leveraging super-resolution images obtained by the so-called Doppler slicing strategy^45^ to search for more apparent differences in vascular features across positions.

Overall, automated brain positioning and neuro-navigation with micro-Doppler images is an issue recently raised by new fUS users, but not yet widely investigated. To date, only one work has addressed the positioning problem through the automated registration of a micro-Doppler scan to a reference^46^. Our approach offers more flexibility as a single image is sufficient to find a position, while CNNs are fast enough for real-time implementation in the neuro-navigation context.

These benefits come at the cost of a need for large datasets to train the classifier. For this proof-of-concept study, we used a standardized dataset with limited sample size and validated the performance on a stroke dataset. Yet, datasets comprising more subjects and more variability (transducer orientation and position, different animal preparation and imaging sequences, further data augmentation, …) will be required to build a model suitable for routine use. In this regard, the increasing adoption of fUS technology by the neuroscience community will facilitate the construction of large databases, especially for mice, the leading mammalian model.

Although our work is validated on anesthetized rats, our strategy is directly applicable to awake imaging and is not conceptually limited to rodents. Micro-Doppler imaging has been successfully applied to humans in neurosurgery^22,47,48^ and non-invasively in newborns by imaging through the fontanel^19,20^. For such clinical applications where accurate positioning is critical, further research will be needed to address the challenges posed by the limited imaging depth and significant differences in vessel scale.

## Methods

### Animals

Experimental procedures were approved by the Committee on Animal Care of the Catholic University of Leuven, in accordance with the national guidelines on the use of laboratory animals and the European Union Directive for animal experiments (2010/63/EU). Adult male Sprague-Dawley rats (n=51; Janvier Labs, France) with an initial weight between 200-300 g were housed in standard ventilated cages and kept in a 12:12 hrs reverse dark/light cycle environment at a temperature of 22 °C with *ad libitum* access to food and water.

### Cranial window for brain-wide imaging and stroke induction

The entire procedure for animal preparation and imaging is performed under isoflurane anesthesia (Iso-Vet, Dechra, Belgium). A mixture of 5% isoflurane in compressed dry air was used to induce anesthesia, subsequently reduced to 2.0-2.5% during surgery and to 1.5% for imaging (see Brunner et al.^43^ for details on surgical procedure). Xylocaine (0.5%, AstraZeneca, England) and Metacam (0.2mg/kg, Boehringer Ingelheim, Canada) were injected subcutaneously as pre-operative and post-operative analgesia, respectively. Intraperitoneal injection of 5% glucose solution was provided every 2 hrs to prevent dehydration. The cranial window extended from Bregma +4.0 to −7.0 mm anteroposterior, laterally ± 6.0 mm was performed in all rats. 26 rats were subjected to stroke by the mean of permanent occlusion of the distal branch of the left middle cerebral artery as detailed in Brunner et al.^43^.

### Micro-Doppler ultrasound imaging of brain vasculature

The data acquisition was performed using an ultrasound imaging scanner equipped with custom acquisition and processing software described in ref.^32^. The scanner is composed of a linear ultrasonic transducer (15 MHz, 128 elements, Xtech15, Vermon, France) connected to 128-channel emission-reception electronics (Vantage, Verasonics, USA) controlled by a high-performance computing workstation (fUSI-2, AUTC, Estonia). The transducer was fixed to a fine-resolution motorized linear stage (T-LSM200A, Zaber Technologies Inc., Canada), allowing for the precise postero-anterior scanning of the brain. The acoustic coupling between the brain and the probe is ensured by a layer of agarose and ultrasound gel (Aquasonic Clear, Parker Laboratories Inc, USA). Each coronal Doppler image is 12.8 mm in width and 9 mm in depth and comprises 300 compound images acquired at 500 Hz. Each compound image is computed by adding nine plane waves (4.5 kHz) with angles from -12° to 12° with a 3° step. The blood signal was extracted from 300 compound images using a single value decomposition filter and removing the 30 first singular vectors^21^. The micro-Doppler image is computed as the mean intensity of the blood signal in these 300 frames, which is an estimator of the cerebral blood volume^13,24^. This sequence enables a temporal resolution of 0.6 sec, an in-plane resolution of 100×110 µm, and an off-plane (image thickness) of 300 µm^32^. Using these parameters, a scan of the brain vasculature consisting of 89 coronal planes spaced by 125 µm was performed between Bregma -6.5 to +3.0 mm.

### Registration of micro-Doppler images

The micro-Doppler 2D scans from all animals were aligned along the anteroposterior axis with respect to recognizable anatomical and vascular patterns (**Fig. 1-c**). This alignment is necessary for correcting potential shifts occurring either during surgery, imaging, or due to inter-animal variability. The process was performed independently by two experts, and any disagreement was resolved post-hoc by consensus. Each micro-Doppler image is then identified by its anatomical position with respect to the Bregma reference point, e.g., Bregma -3.0 mm.

### Datasets generation

Several datasets have been extracted from the initial scans using a down-sampling factor ranging from 2 to 5. This corresponds to an artificial increase in the spacing between two consecutive micro-Doppler images. To create the dataset associated with a given factor F, we extracted images from Bregma -3.0 mm with a spacing of Fx125 µm, within the limits of the cranial windows, thus resulting in 77 common positions across animals (**Fig. 1-b**). The 5 datasets spaced by [250, 375, 500, 625, 750] µm, respectively, contain [39, 26, 20, 15, 13] different positions. 50% of the animals were randomly selected for training, 25% for tuning the hyperparameters (i.e., validation), and the remaining 25% for evaluating the final performances of the model (i.e., test).

### Image preprocessing

To increase the contrast, a correction factor (power of 0.25) has been applied to every pixel of all images in each dataset. The intensity amplitude was then normalized to fit into a [0, 1] interval. The overall process has been implemented using MATLAB (R2018b, Mathworks, USA).

### Model selection

To select the optimal model for the experiments, we evaluated 5 convolutional neural network architectures (CNN; ResNet50^49^, DenseNet121^39^, VGG11^50^, ViT-Base-16^51^, and EfficientNetV2-S^52^), support vector machines (SVM) with different feature extraction methods (HOG^40^, SIFT^53^, PCA) and kernels^54^. These models were selected for their compatibility with datasets of relatively small sample sizes. Both ResNet50-CNN and DenseNet121-CNN were pretrained on ImageNet^55^ as previously suggested^56^. The default parameters and architectures were used, as implemented in the ‘torchvision’ (PyTorch, version 0.7.0) Python package. The network’s last layer -the classifier -was replaced by a fully connected layer outputting n values, n being the number of positions in the training set, and passed through a softmax layer afterward. Both CNN and SVM models were trained and evaluated on the dataset with 375 µm spacing, corresponding to the smallest spacing above the anteroposterior spatial resolution of the modality employed. It, therefore, corresponds to the largest dataset without overlapping information. SVMs were implemented using the ‘scikit-learn’ (version 0.23.1) Python package.

### Training and evaluation procedures for CNNs

For each of the datasets used in this work, images were resized to 224×320 pixels by bicubic interpolation, and their grey channel extended in RGB to fit the ImageNet format imposed by the pre-training^55^. All the data were normalized with the mean and standard deviation of the full dataset. We augmented the size of the training set with rotations of ±4°and ±8° applied to all micro-Doppler images. The weights of the network were optimized with the Adam algorithm using a cross-entropy loss function. The hyperparameters were selected through a random search (see **Supplementary Table S3** and **S4**). Other parameters were kept with default values. The final model performance was then evaluated on the testing set. The overall procedure has been performed on a single computer equipped with bi-Xeon E5-2620 CPU (Intel, USA), 64 GB RAM, and 4 RTX2080 (8 GB) GPUs (Nvidia, USA).

### Probability-based regression

This step occurs after the training of the model for the classification task. When an input image is fed to the network, it outputs a probability *P*(*K*_i_) for each one of the positions (*K*_i_)_1≤*i*≤*N*_ included in the training set. We use these probabilities to obtain an estimate 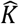 of the position *K* using the following equation:

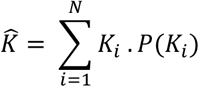

The positioning error is defined as 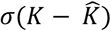, where σ denotes the standard deviation.

### Visualization of relevant features for image classification using GradCAM

We extracted the pixels in the input image, driving the classification using the Gradient-weighted Class Activation Map (GradCAM) technique, following the recommendations of Adebayo *et al*.^57^ on the relevant visualization approaches. This method aggregates the gradients associated with the prediction for each feature map in a given layer to produce a coefficient measuring the contribution of each of the maps to the network’s prediction. Here, the gradients and feature maps were extracted at the last layer before the classifier. The output heatmaps were then resized by bilinear interpolation to the original image, and a threshold of 0.7 was applied to limit the effect of the interpolation.

### GradCAM registration on the atlas for anatomical extraction

We used a digital version of the rat brain Paxinos atlas^6,43^ to extract the anatomical regions associated with the GradCAM heatmaps. The input scan was taken as a volume and interpolated to fit the atlas resolution (50×50×50 µm^3^ voxel size). A 3D rigid registration was performed using a MATLAB custom script^32,43^. This procedure has been applied to all the samples from the validation and testing sets by an expert. To extract the regions from the GradCAM heatmap, a volume (89 planes as the input data) was constructed from the heatmaps by zero-padding the missing sections before applying the transformation matrix.

### Evaluation of the stroke dataset

We used 26 rats subjected to stroke (see above and ref.^43^). All rats were imaged in the original dataset, and 14/7/7 individuals were respectively present in training/validation/testing sets. The scans were registered, and a dataset with 375-µm spacing was created following the same procedure as for the previous experiment. The class predictions were obtained by processing the images through the DenseNet121-CNN previously trained on the pre-stroke dataset with 375-µm spacing without re-training.

## Supplementary Materials

**Supplementary Fig. S1:**
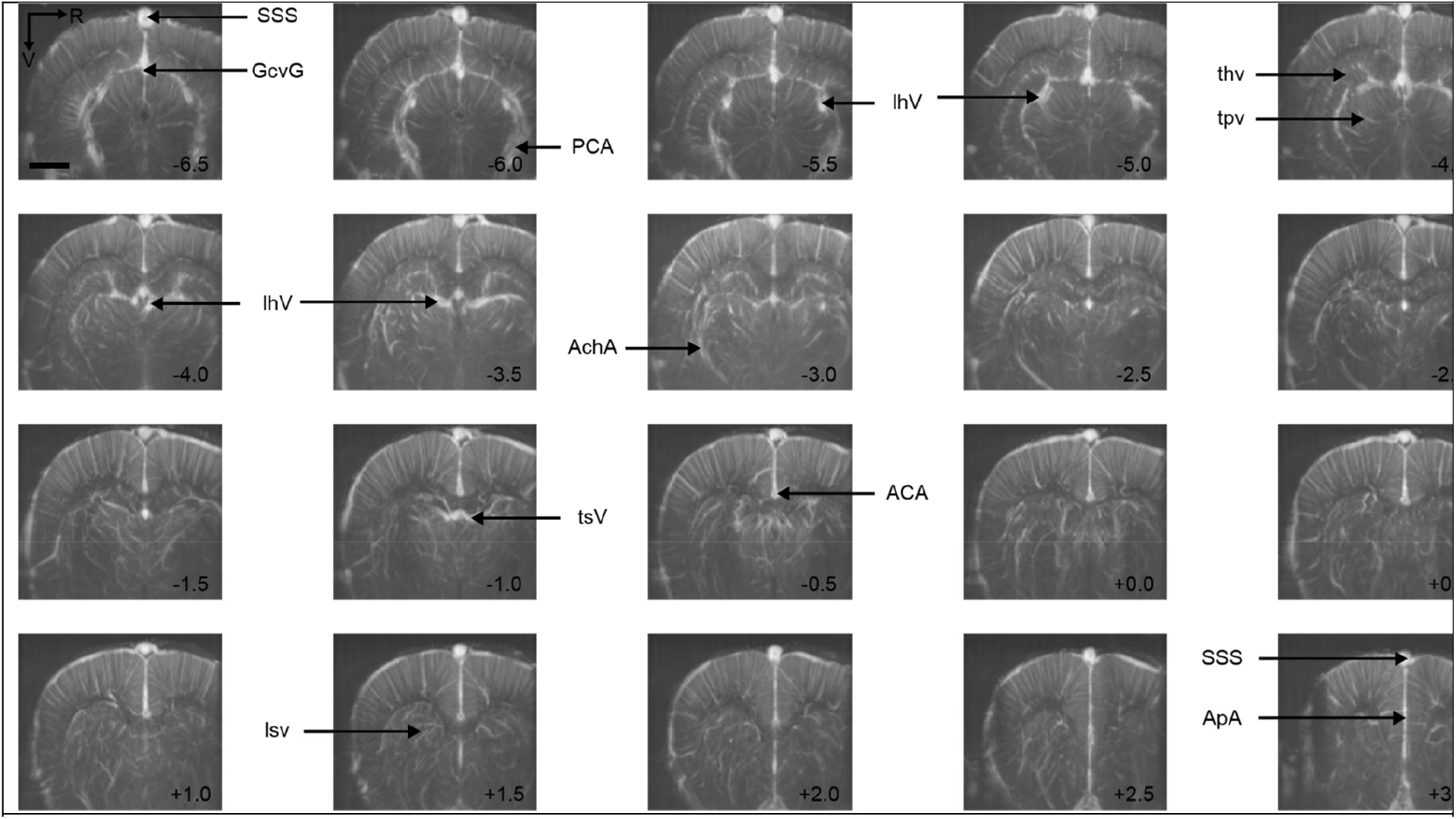
Micro-Doppler images extracted from the dataset 500_20_ covering a large part of the rat brain from posterior (top left) to anterior positions (bottom right). Major vessels are identified as follow: ACA: anterior cerebral artery, AchA: anterior choroidal artery, ApA: azygos pericallosal artery, GcvG: great cerebral vein of Galen, lhV: longitudinal hippocampal vein, lsv: lenticulostriate vessels, thv: transverse hippocampal vessels, tsV: thalamostriate vein, tpv: thalamoperforating vessels, SSS: superior sagittal sinus, A: anterior, L: left, V: ventral. The Bregma position (in mm) of the micro-Doppler image is given on the lower right corner. Scale bar: 2 mm.

**Supplementary Fig. S2.**
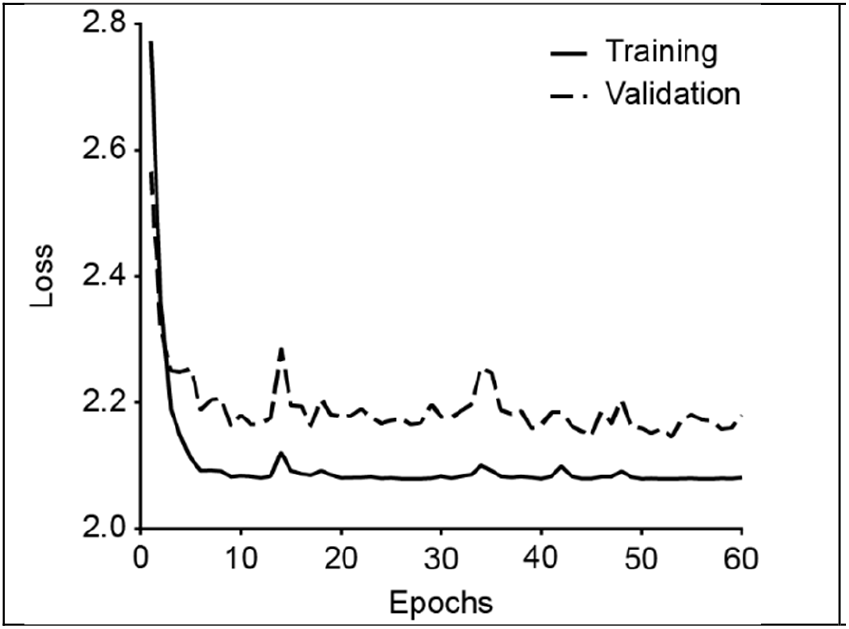
Evolution of the cross-entropy loss with epochs on dataset 500_20_ during training (plain line) and validation (dashed line).

**Supplementary Table S1.** Performance evaluation of selected models on dataset with 375-µm spacing.

| Model | Validation accuracy (%) | Testing accuracy (%) | Positioning error (μm) |
| --- | --- | --- | --- |
| ResNet50-CNN | 82.5 | 77.2 | 313 |
| **DenseNet121-CNN (main)** | **85.2** | **81.9** | **176** |
| VGG11-CNN | 81.9 | 77.8 | 327 |
| ViT-Base-16-CNN | 81.1 | 80.8 | 178 |
| EfficientNetV2-S-CNN | 81.7 | 80.5 | 176 |
| HOG-SVM $\chi^2$ (baseline) | 71.6 | 80.4 | / |
| HOG-SVM rbf | 65.1 | 68.3 | / |
| SIFT-SVM rbf | 29.3 | 24.5 | / |
| PCA-SVM rbf | 53.5 | 64.5 | / |

**Supplementary Table S2.** DenseNet121-CNN and HOG-SVM performance metrics on the datasets with different spacings. ^*^p-value<0.05, ^**^p-value<0.01.

| Spacing (μm) | | 250 | 375 | 500 | 625 | 750 |
| --- | --- | --- | --- | --- | --- | --- |
| Number of positions |  | 39 | 26 | 20 | 15 | 13 |
| **DenseNet121-CNN** | Validation accuracy (%) | 64.3 | 85.2 | 95.4 | 100 | 100 |
|  | Test accuracy (%) | 56.2 | 81.9 | 93.1 | 96.9 | 98.2 |
|  | Difference between test and validation accuracy | 8.1 | 3.3 | 2.3 | 3.1 | 1.8 |
|  | Maximum class accuracy test (%) | 84.6 | 100 | 100 | 100 | 100 |
|  | Minimum class accuracy test (%) | 0.0 | 30.8 | 84.6 | 84.6 | 84.6 |
| **HOG-SVM** | Validation accuracy (%) | 54 | 71.6 | 81.9 | 88.7 | 92.9 |
|  | Test accuracy (%) | 58 | 80.4 | 85 | 90.8 | 95.9 |
|  | Difference between test and val. Accuracy | -4 | -8.8 | -3.1 | -2.1 | -3 |
|  | Maximum class accuracy (%) | 84.6 | 100 | 100 | 100 | 100 |
|  | Minimum class accuracy (%) | 20.3 | 61.5 | 53.8 | 53.8 | 84.6 |
| **McNemar test** | Test statistic | 86.000 | 28.000 | 12.000 | 4.000 | 1.000 |
|  | P-value | 0.552 | 0.450 | 0.002 (**) | 0.012 (*) | 0.219 |

**Supplementary Table S3.** Hyperparameters used for training CNNs on dataset with 375-µm spacing.

| Model | Learning rate | Weight decay | Batch size | Epochs |
| --- | --- | --- | --- | --- |
| ResNet50-CNN | 0.00016611147361060297 | 6.597427678401722e-05 | 64 | 30 |
| DenseNet121-CNN (main) | 0.00019968374184068966 | 1.0149428174533185e-05 | 32 | 60 |
| VGG11-CNN | 5.489552742460815e-05 | 0.004436151585657852 | 64 | 30 |
| ViT-Base-16-CNN | 7.910139901190354e-05 | 1.8030985758135736e-06 | 64 | 30 |
| EfficientNetV2-S-CNN | 1.6852664243005598e-05 | 0.0002395035729247215 | 32 | 30 |

**Supplementary Table S4.**
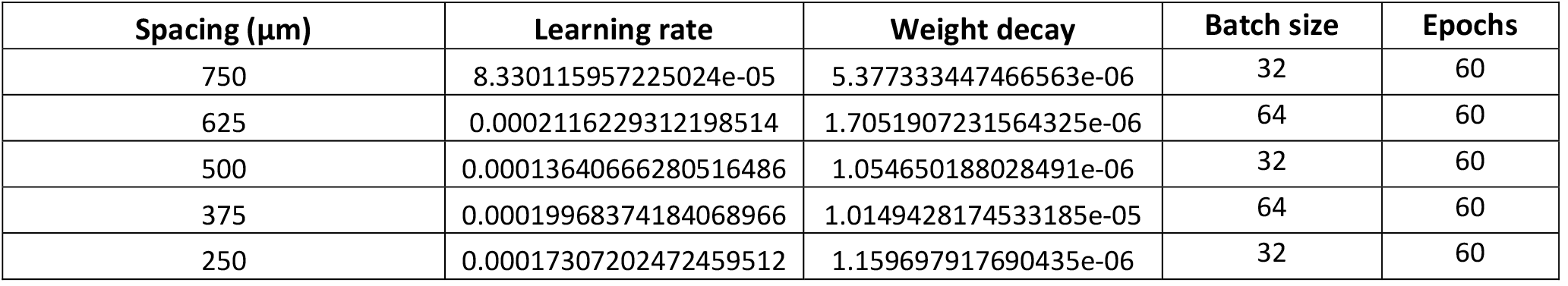
Hyperparameters used for training DenseNet121-CNN model on dataset per spacing.

| Spacing (μm) | Learning rate | Weight decay | Batch size | Epochs |
| --- | --- | --- | --- | --- |
| 750 | 8.330115957225024e-05 | 5.377333447466563e-06 | 32 | 60 |
| 625 | 0.0002116229312198514 | 1.7051907231564325e-06 | 64 | 60 |
| 500 | 0.00013640666280516486 | 1.054650188028491e-06 | 32 | 60 |
| 375 | 0.00019968374184068966 | 1.0149428174533185e-05 | 64 | 60 |
| 250 | 0.00017307202472459512 | 1.159697917690435e-06 | 32 | 60 |

## Acknowledgments

This work was supported by grants from Europe: ERANET-NEURON (G0G3721N-UNSCRAMBLY) and MSCA training network (SOPRANI-GAP-101119916), from Belgium: FWO senior research grants (G079623N-TRPM3 and G0C9923N-USNI4TBI) and FWO senior postdoctoral fellowship (12D7523N-BRAIN PAIN) and from USA: NIH (R01NS129836-01A1). This research also received funding from IMEC and the Flemish regional government (AI Research Program). We thank the NERF animal caretakers including I. Eyckmans, F. Ooms, and S. Luijten, for their help managing the animals.

We extend our gratitude to Dr. Matthew Blaschko and Dr. Johan Suykens for their valuable input on the manuscript and overall guidance in overseeing T.L.’s Ph.D. training.

## Author contributions

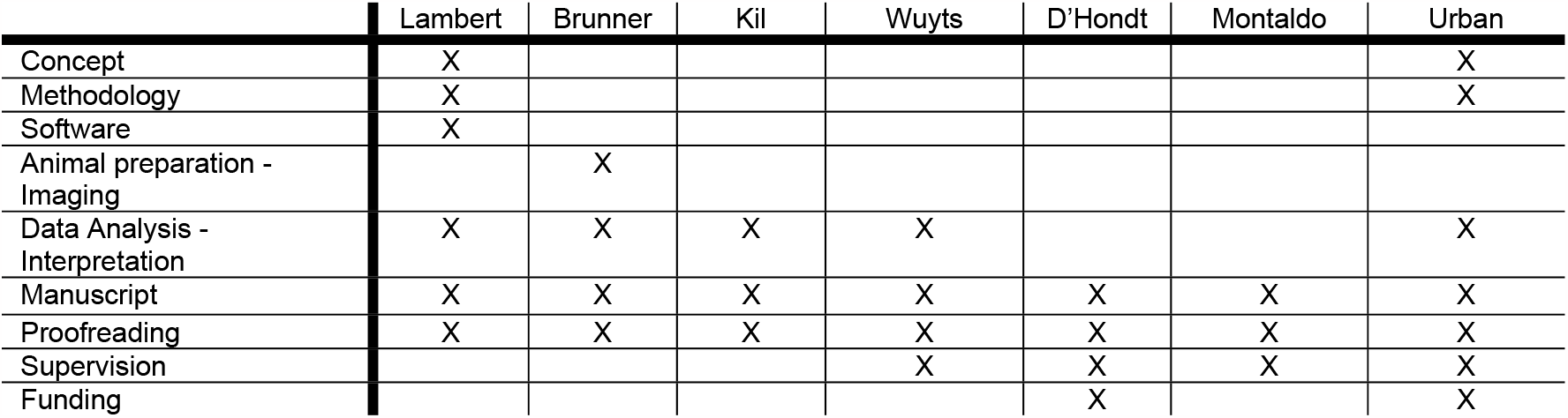

## References

1. Enchev, Y. Neuronavigation: geneology, reality, and prospects. Neurosurg. Focus 27, E11 (2009).

2. Thomas, N. W. D. & Sinclair, J. Image-guided neurosurgery: History and current clinical applications. J. Med. Imaging Radiat. Sci. 46, 331–342 (2015).

3. Wang, M. N. & Song, Z. J. Classification and analysis of the errors in neuronavigation. Neurosurgery 68, 1131–43; discussion 1143 (2011).

4. Alam, F., Rahman, S. U., Ullah, S. & Gulati, K. Medical image registration in image guided surgery: Issues, challenges and research opportunities. Biocybern. Biomed. Eng. 38, 71–89 (2018).

5. Wang, Q. et al. The Allen mouse brain common coordinate framework: A 3D reference atlas. Cell 181, 936–953.e20 (2020).

6. Paxinos, G. & Watson, C. Rat brain in stereotaxic coordinates. (Academic Press, 2007).

7. Niedworok, C. J. et al. aMAP is a validated pipeline for registration and segmentation of high-resolution mouse brain data. Nat. Commun. 7, 11879 (2016).

8. Fuglstad, J. G., Saldanha, P., Paglia, J. & Whitlock, J. R. HERBS: Histological E-data Registration in rodent Brain Spaces. bioRxiv (2021) doi:10.1101/2021.10.01.462770.

9. Xiao, D., Forys, B. J., Vanni, M. P. & Murphy, T. H. MesoNet allows automated scaling and segmentation of mouse mesoscale cortical maps using machine learning. Nat. Commun. 12, 5992 (2021).

10. Qu, L. et al. Cross-modal coherent registration of whole mouse brains. Nat. Methods 19, 111–118 (2022).

11. Montijn, J. S. & Heimel, J. A. A universal pipeline for the alignment of electrode tracks with slice histology and electrophysiological data. bioRxiv (2022) doi:10.1101/2022.06.20.496782.

12. Giovannucci, A. et al. CaImAn an open source tool for scalable calcium imaging data analysis. Elife 8, (2019).

13. Macé, E. et al. Functional ultrasound imaging of the brain. Nat. Methods 8, 662–664 (2011).

14. Urban, A. et al. Real-time imaging of brain activity in freely moving rats using functional ultrasound. Nature methods 12, 873–878 (2015).

15. Macé, É. et al. Whole-brain functional ultrasound imaging reveals brain modules for visuomotor integration. Neuron 100, 1241–1251.e7 (2018).

16. Brunner, C. et al. A platform for brain-wide volumetric functional ultrasound imaging and analysis of circuit dynamics in awake mice. Neuron 108, 861–875.e7 (2020).

17. Dizeux, A. et al. Functional ultrasound imaging of the brain reveals propagation of task-related brain activity in behaving primates. Nat. Commun. 10, 1400 (2019).

18. Takahashi, D. Y. et al. Social-vocal brain networks in a non-human primate. BioRxiv (2021) doi:10.1101/2021.12.01.470701.

19. Demene, C. et al. Functional ultrasound imaging of the brain activity in human neonates. in 2016 IEEE International Ultrasonics Symposium (IUS) (IEEE, 2016). doi:10.1109/ultsym.2016.7728657.

20. Demene, C. et al. Functional ultrasound imaging of brain activity in human newborns. Sci. Transl. Med. 9,(2017).

21. Urban, A. et al. Functional Ultrasound Imaging of Cerebral Capillaries in Rodents and Humans. Jacobs Journal of Molecular and Translational Medicine 1, (2015).

22. Soloukey, S. et al. Functional Ultrasound (fUS) During Awake Brain Surgery: The Clinical Potential of Intra-Operative Functional and Vascular Brain Mapping. Front. Neurosci. 13,(2020).

23. Soloukey, S. et al. High-resolution micro-Doppler imaging during neurosurgical resection of an arteriovenous malformation: illustrative case. J Neurosurg Case Lessons 4,(2022).

24. Mace, E. et al. Functional ultrasound imaging of the brain: theory and basic principles. IEEE Trans. Ultrason. Ferroelectr. Freq. Control 60, 492–506 (2013).

25. Edelman, B. J. & Macé, E. Functional ultrasound brain imaging: Bridging networks, neurons, and behavior. Curr. Opin. Biomed. Eng. 18, 100286 (2021).

26. Montaldo, G., Urban, A. & Macé, E. Functional ultrasound neuroimaging. Annu. Rev. Neurosci. 45, 491–513 (2022).

27. Bercoff, J. et al. Ultrafast compound Doppler imaging: providing full blood flow characterization. IEEE Trans. Ultrason. Ferroelectr. Freq. Control 58, 134–147 (2011).

28. Urban, A. et al. Chronic assessment of cerebral hemodynamics during rat forepaw electrical stimulation using functional ultrasound imaging. Neuroimage 101, 138–149 (2014).

29. Sans-Dublanc, A. et al. Optogenetic fUSI for brain-wide mapping of neural activity mediating collicular-dependent behaviors. Neuron 109, 1888–1905.e10 (2021).

30. Nunez-Elizalde, A. O. et al. Neural correlates of blood flow measured by ultrasound. Neuron 110, 1631–1640.e4 (2022).

31. Sieu, L.-A. et al. EEG and functional ultrasound imaging in mobile rats. Nat. Methods 12, 831–834 (2015).

32. Brunner, C. et al. Whole-brain functional ultrasound imaging in awake head-fixed mice. Nat. Protoc. 16, 3547–3571 (2021).

33. Richtsmeier, J. T., Baxter, L. L. & Reeves, R. H. Parallels of craniofacial maldevelopment in Down syndrome and Ts65Dn mice. Dev. Dyn. 217, 137–145 (2000).

34. Xiao, J. A new coordinate system for rodent brain and variability in the brain weights and dimensions of different ages in the naked mole-rat. J. Neurosci. Methods 162, 162–170 (2007).

35. Dorr, A., Sled, J. G. & Kabani, N. Three-dimensional cerebral vasculature of the CBA mouse brain: a magnetic resonance imaging and micro computed tomography study. Neuroimage 35, 1409–1423 (2007).

36. Xiong, B. et al. Precise cerebral vascular atlas in stereotaxic coordinates of whole mouse brain. Front. Neuroanat. 11, 128 (2017).

37. Todorov, M. I. et al. Machine learning analysis of whole mouse brain vasculature. Nat. Methods 17, 442–449 (2020).

38. Nouhoum, M. et al. A functional ultrasound brain GPS for automatic vascular-based neuronavigation. Sci. Rep. 11, 15197 (2021).

39. Huang, G., Liu, Z., Van Der Maaten, L. & Weinberger, K. Q. Densely connected convolutional networks. in 2017 IEEE Conference on Computer Vision and Pattern Recognition (CVPR) (IEEE, 2017). doi:10.1109/cvpr.2017.243.

40. Dalal, N. & Triggs, B. Histograms of oriented gradients for human detection. in 2005 IEEE Computer Society Conference on Computer Vision and Pattern Recognition (CVPR’05) (IEEE, 2005). doi:10.1109/cvpr.2005.177.

41. McNEMAR, Q. Note on the sampling error of the difference between correlated proportions or percentages. Psychometrika 12, 153–157 (1947).

42. Selvaraju, R. R. et al. Grad-CAM: Visual explanations from deep networks via gradientbased localization. in 2017 IEEE International Conference on Computer Vision (ICCV) (IEEE, 2017). doi:10.1109/iccv.2017.74.

43. Brunner, C. et al. Brain-wide continuous functional ultrasound imaging for real-time monitoring of hemodynamics during ischemic stroke. (2022) doi:10.1101/2022.01.19.476904.

44. Geirhos, R. et al. ImageNet-trained CNNs are biased towards texture; increasing shape bias improves accuracy and robustness. arXiv [cs.CV] (2018).

45. Bar-Zion, A. et al. Doppler slicing for ultrasound super-resolution without contrast agents. bioRxiv (2021) doi:10.1101/2021.11.19.469083.

46. Nouhoum, M. et al. Fully-automatic ultrasound-based neuro-navigation : The functional ultrasound brain GPS. Research Square (2021) doi:10.21203/rs.3.rs-382732/v1.

47. Imbault, M. et al. Functional ultrasound imaging of the human brain activity: An intraoperative pilot study for cortical functional mapping. in 2016 IEEE International Ultrasonics Symposium (IUS) (IEEE, 2016). doi:10.1109/ultsym.2016.7728505.

48. Imbault, M., Chauvet, D., Gennisson, J.-L., Capelle, L. & Tanter, M. Intraoperative Functional Ultrasound Imaging of Human Brain Activity. Sci. Rep. 7, 7304 (2017).

49. He, K., Zhang, X., Ren, S. & Sun, J. Deep Residual Learning for Image Recognition. in 2016 IEEE Conference on Computer Vision and Pattern Recognition (CVPR) 770–778 (IEEE, 2016).

50. Simonyan, K. & Zisserman, A. Very deep convolutional networks for large-scale image recognition. arXiv [cs.CV] (2014).

51. Dosovitskiy, A. et al. An image is worth 16x16 words: Transformers for image recognition at scale. arXiv [cs.CV] (2020).

52. Tan, M. & Le, Q. V. EfficientNetV2: Smaller models and faster training. arXiv [cs.CV] (2021).

53. Lowe, D. G. Distinctive image features from scale-invariant keypoints. Int. J. Comput. Vis. 60, 91–110 (2004).

54. Zhang, J., Marszałek, M., Lazebnik, S. & Schmid, C. Local features and kernels for classification of texture and object categories: A comprehensive study. Int. J. Comput. Vis. 73, 213–238 (2007).

55. Deng, J. et al. ImageNet: A large-scale hierarchical image database. in 2009 IEEE Conference on Computer Vision and Pattern Recognition (IEEE, 2009). doi:10.1109/cvpr.2009.5206848.

56. Tajbakhsh, N. et al. Convolutional neural networks for medical image analysis: Full training or fine tuning? IEEE Trans. Med. Imaging 35, 1299–1312 (2016).

57. Adebayo, J. et al. Sanity Checks for Saliency Maps. arXiv [cs.CV] (2018).

